# Electrical brain activations in preadolescents during a probabilistic reward-learning task reflect cognitive processes and behavioral strategy

**DOI:** 10.1101/2023.10.16.562326

**Authors:** Yu Sun Chung, Berry van den Berg, Kenneth C. Roberts, Armen Bagdasarov, Marty G. Woldorff, Michael S. Gaffrey

**Affiliations:** Department of Psychology and Neuroscience, Duke University, Reuben-Cooke Building, 417 Chapel Drive, Durham, NC 27708, USA; Department of Psychology, Kean University, East Campus, 1000 Morris Avenue, Union, NJ, 07083, USA; Experimental Psychology, University of Groningen, The Netherlands; Center for Cognitive Neuroscience, Department of Psychiatry, Psychology & Neuroscience and Neurobiology, Duke University, Durham, NC, 27708 USA; Children’s Wisconsin, 9000 W. Wisconsin Avenue, Milwaukee, WI, 53226; Medical College of Wisconsin, Division of Pediatric Psychology and Developmental Medicine, Department of Pediatrics, 8701 Watertown Plank Road, Milwaukee, WI, 53226

**Keywords:** reinforcement learning, reward-related positivity, attention, N2pc, P300, win-stay-lose-switch strategy

## Abstract

Both adults and children learn through feedback which environmental events and choices are associated with higher probability of reward, an ability thought to be supported by the development of fronto-striatal reward circuits. Recent developmental studies have applied computational models of reward learning to investigate such learning in children. However, tasks and measures effective for assaying the cascade of reward-learning neural processes in children have been limited. Using a child-version of a probabilistic reward-learning task while recording event-related-potential (ERP) measures of electrical brain activity, this study examined key processes of reward learning in preadolescents (8-12 years old; n=30), namely: (1) reward-feedback sensitivity, as measured by the early-latency, reward-related, frontal ERP positivity, (2) rapid attentional shifting of processing toward favored visual stimuli, as measured by the N2pc component, and (3) longer-latency attention-related responses to reward feedback as a function of behavioral strategies (i.e., Win-Stay-Lose-Shift), as measured by the central-parietal P300. Consistent with our prior work in adults, the behavioral findings indicate preadolescents can learn stimulus-reward outcome associations, but at varying levels of performance. Neurally, poor preadolescent learners (those with slower learning rates) showed greater reward-related positivity amplitudes relative to good learners, suggesting greater reward-feedback sensitivity. We also found attention shifting towards to-be-chosen stimuli, as evidenced by the N2pc, but not to more highly rewarded stimuli as we have observed in adults. Lastly, we found the behavioral learning strategy (i.e., Win-Stay-Lose-Shift) reflected by the feedback-elicited parietal P300. These findings provide novel insights into the key neural processes underlying reinforcement learning in preadolescents.

## 1. Introduction

From an early age, children learn about the world around them through trial and error. That is, through repeated experience, children progressively learn what actions are likely to result in relatively positive or negative outcomes given a specific environment and/or set of choices. The ability to learn associations between choice behavior and outcomes and to then change patterns of subsequent choice behavior accordingly, has been called probabilistic reinforcement learning (PRL). Computational models of PRL often separate such learning into computationally-driven components associated with both behaviorally and neurobiologically distinguishable processes. Specifically, PRL models involve a sequence of hypothesized neurocognitive processes, including evaluation of the relative value associated with available choices, updating of stimulus-reward outcome associations based on feedback, and shaping of selective attention toward a more-likely-winning stimulus to guide future choice (Anderson, 2017). However, while some of the neural correlates of PRL have been well studied in adults, relatively few studies have investigated this question in adolescents or children (reviewed by (DePasque & Galván, 2017). This is a critical knowledge of gap because the study of PRL in children could be quite valuable for identifying specific neurocognitive processes that may account for differences in these important cognitive functions across development (Nussenbaum & Hartley, 2019b) and may also predict the risk for depression, which is heightened during adolescence (Chen et al., 2015; Keren et al., 2018).

At the behavioral level, PRL is evident during childhood, adolescence, and adulthood (Raab & Hartley, 2018). However, findings from studies using computational models of PRL to examine how value-based learning ability changes with age have provided mixed results. As reviewed by (Nussenbaum & Hartley, 2019), some developmental studies have reported that learning rates increase from childhood to adulthood (Davidow et al., 2016; Master et al., 2020) while others have reported that they do not change with age (Palminteri et al., 2016). Such mixed behavioral findings may reflect individual differences in neurobiological alterations in one or all of the processes underlying PRL (i.e., value representation associated with choices, updating of stimulus-reward outcome associations, shaping attention towards reward-predictive cues). For example, learning rates captures individual differences in the value-updating process, reflecting the degree to which reward feedback and reward prediction errors are incorporated into the updated value estimates during learning. However, existing studies in children rarely considered individual learning rates in the investigation of key neural components of reinforcement learning. A number of empirical studies using functional Magnetic Resonance Imaging (fMRI) in adults have suggested that reward sensitivity is signaled by dopaminergic midbrain-fronto-striatal reward brain circuits (Averbeck & O’Doherty, 2022; Nasser et al., 2017; Schultz, 2016), including in adolescents (Christakou et al., 2013; Cohen et al., 2010; Hauser et al., 2015). However, the sluggishness of fMRI signals limits the ability of this method to elucidate the cascade of the different subprocesses underlying learning on a trial-by-trial basis, which requires more precise temporal resolution.

The high temporal resolution of electroencephalogram (EEG) measures of brain activity makes such recordings particularly well-suited for studying the neural subprocesses associated with PRL. More specifically, the ability to extract and measure event-related potentials (ERPs) time-locked to stimulus and reward-feedback events provides a particularly effective opportunity to delineate the temporal cascade of neurocognitive processes related to reward processing within a trial (Amodio et al., 2014). One ERP component that has been extensively examined in reward studies is the reward positivity (RewP), a frontal, positive-polarity, brain wave measured as the response to gain feedback relative to that for loss feedback, occurring approximately 250-350ms after that feedback. With respect to RL theory (Holroyd & Coles, 2002; Sutton & Barto, 2018), the RewP is thought to index reward-related sensitivity in the mesocortical dopamine system, specifically reflecting reward prediction error signal that is sensitive to outcome valence and is larger for unexpected positive events relative to unexpected negative events (Holroyd & Coles, 2002). Although the RewP (or its inverse, the Feedback-related Negativity, FN, calculated as loss-minus gain feedback in some earlier research (Krigolson, 2018)) as a measure of response to reward has been widely investigated in adults (see (Glazer et al., 2018; Kujawa, 2024) for reviews), its role in PRL during this age has not. For example, we know little about how individual learning rates in adults or preadolescents may modulate changes in the RewP amplitudes, and other key neural components of PRL, as described in the below.

As one learns which stimuli are more likely to generate reward outcomes, one tends to pay more attention to those stimuli. Such attention shifting toward reward-associated stimuli has been indexed by the lateralized attention-shifting-sensitive N2pc component in prior work with adults (Hickey et al., 2010; San Martin et al., 2016;). In contrast, there have been only a handful of ERP studies that have examined the N2pc in children, either in the context of a reward task or more generally during visual search (Couperus & Quirk, 2015; Li et al., 2022; Shimi et al., 2014; Sun et al., 2018; Turoman et al., 2021) and/or working memory tasks (Rodríguez-Martínez et al., 2021; Shimi et al., 2015). Consistent with prior work in adults, existing studies with children indicate the elicitation of the N2pc component during non-PRL attentional tasks, indicating that this component can be used as a marker of attentional selection in children as it is with adults (Couperus & Quirk, 2015; Li et al., 2022; Shimi et al., 2014; Sun et al., 2018; Turoman et al., 2021). However, the presence of this attentional neural process has not been directly tested within the framework of computational PRL models. Accordingly, we know little about the functionality of the N2pc ERP component in children within a PRL framework – that is, whether it may reflect an index of early attentional shifting as a function of reward learning in children, as has been shown previously in adults.

Lastly, the P300 has been thought more generally to reflect longer-latency attentional processes involved in information updating of salient outcomes during evaluative processing (for reviews, see <u>Glazer et al., 2018</u>, <u>San Martín, 2012</u>). The P300 is a centro-parietally distributed, positive-going wave that peaks between 400 and 600 ms following a stimulus (San Martín et al., 2013; Wu & Zhou, 2009). It has been successfully used to investigate PRL with adults (e.g., (Donaldson et al., 2016). More specifically, prior work has shown that P300 is sensitive to a longer-latency top-down-control process of outcome evaluation, suggesting that outcome-related feedback information in a probability choice task (e.g., probability and magnitude) is used to guide future decisions for the goal of reward maximization (<u>San Martín et al., 2013</u>, <u>Wu and Zhou, 2009</u>, <u>Zhang et</u> <u>al., 2013</u>). However, like the N2pc, the functionality of the feedback locked P300 (reward gain vs. loss feedback) during PRL in children has been little examined.

There were several main goals of the current study. First, we wanted to establish the feasibility and initial validity of a developmentally appropriate probabilistic reward learning task, which we have termed the C-PLearn task (for Child Probability Learning task). Based on our results, we believe that this task would be a promising tool for studying both typical and atypical development of key neurocognitive processes underlying PRL. For example, alterations in PRL abilities have been known to be critical components of hedonic functioning that predicts depression risk for children and adolescents (Keren et al., 2018; Morris et al., 2015; Saulnier et al., 2023). The design of the C-PLearn task was based on our prior work investigating neural correlates of PRL processes in adults (van den Berg et al., 2019). Secondly, we wanted to use this task to examine key putative subprocesses of PRL in the instantiation of heuristic behavioral learning strategies, namely, Win-Stay-Lose-Shift (WSLS) (Hernstein et al., 2000), in a group of preadolescent children for the first time: namely the subprocesses of (1) reward-feedback sensitivity, as measured by the RewP, (2) early attentional processing toward stimuli with higher reward value or reward likelihood as a shaped by reward learning, as measured by the N2pc, (3) longer-latency attentional processing in response to the feedback as a function of behavioral learning strategies (i.e., Win-Stay-Lose-Shift, WSLS (Hernstein et al., 2000)), as measured by the central-parietal P300. More specifically, we wanted to assay the RewP (reflecting reward-feedback sensitivity), the N2pc (reflecting the shifting of attention to certain stimuli versus others), and the P300 (reflecting longer-latency attention-related cognitive processes) as a function of reward learning for preadolescents within the PRL framework. In terms of hypotheses, we predicted that preadolescents who didn’t learn the stimulus-outcome associations very well (i.e., “bad” learners) would show smaller RewP amplitudes, consistent with the RL framework, compared to “good” learners, and/or would adopt less effective behavioral strategies. Finally, we hypothesized that the central-parietal, feedback-locked P300 may reflect a modulation of longer-latency cognitive processing in the updating of reward value as a function of behavior strategy for preadolescent PRL learning (e.g., win-stay-lose-switch), an idea that has been supported by the literature in adults (Donaldson et al., 2016; von Borries et al., 2013).

## 2. Methods

### 2.1. Participants

Thirty children between 8 and 12 years old (mean age = 9.94 years ±1.26 (SD); 15 female; 30 right-handed; 20% Hispanic or Latino ethnicity, 80% non-Hispanic white) were recruited from a Research Participation Database at Duke University maintained by the Human Subjects Coordinator for the Department of Psychology and Neuroscience. In addition, methods included posting flyers on social media sites and distributing flyers to local schools were used to recruit participants.

Children who were color blind, reported a current and/or historical diagnosis of any psychiatric disorder, had a history of neurological disorder or insult (e.g., seizures, hydrocephalus, cerebral palsy, brain tumor, extended loss of consciousness, head trauma), took psychotropic mediation for a mood or behavioral difficulty, or were left-handed were excluded from study participation. At the end of the EEG sessions, a $30 Amazon e-gift card was sent to the parents via email for the participation of their children in the study. Children also received candy and/or small toys after each completed session. Additional compensation was provided ($10 Amazon e-gift card per hour) to parents if testing sessions exceeded 2.5 hours. This study was conducted in accordance with protocols approved by the Duke Institutional Review Board.

### 2.2. The Child Probabilistic Reward Learning (C-PLearn) and Localizer Tasks

The C-PLearn task used in the current study for children was based on our adult PLearn task previously used successfully in college students (van den Berg et al., 2019). Participants completed the C-PLearn, followed by a face-vs-house neural localizer task (for localizing the face-selective neural activity on the scalp), while EEG was recorded. The task was programmed and run on the OpenSesame platform for behavioral research (Mathôt et al., 2012), and event codes were sent to the EEG acquisition computer using the Python ‘egi’ package.

C-PLearn: C-PLearn consisted of 20 blocks of 18 trials each, slightly less than the 20 trials per block in the original adult version to accommodate reduced attention spans and lower tolerance for prolonged experimental tasks in children. Also, the probability of reward was set to 0.77 (instead of ranging randomly from 0.5 to 0.75 as in the original adult version) to mitigate frustration in the children and potential confusion noted during early piloting sessions using the original adult reward probability levels. At the beginning of each trial in a block, participants were presented with two images on the screen, one of a face and one of a house, and were instructed to choose between the two (Figure 1). After the participants made their choice on each trial, they received feedback as to whether they won points or lost points on that trial. Participants were instructed to try to learn during each block whether faces or houses were more likely to result in winning points, and that winning more points would allow them to have more points for choosing small toys that had differing levels of cost. At the end of each 18-trial block, feedback was given as to how many points the participant had accrued up until that point. After a practice period in which the subject needed to demonstrate understanding of how to do the task, the test session started with the C-PLearn task runs.

**Figure 1.**
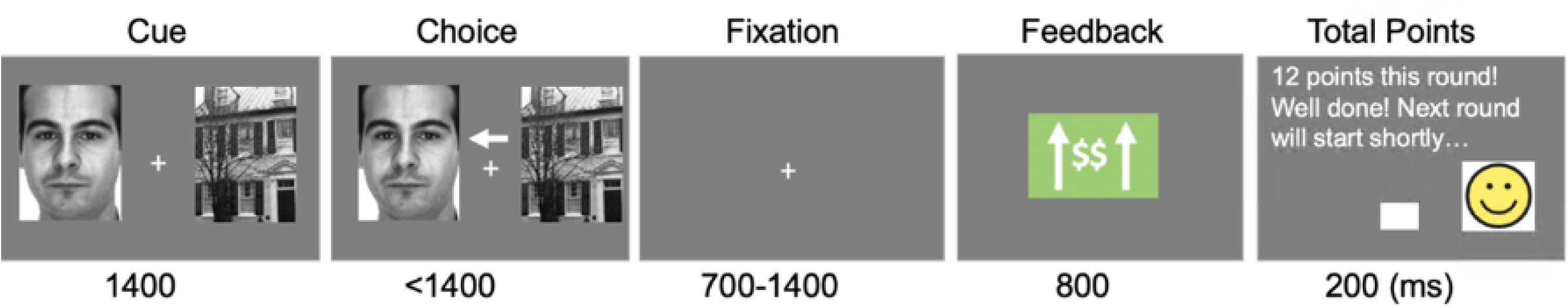
Child-version of Probabilistic Reward Learning (C-PLearn) Task. An example of one trial with reward feedback (i.e., green box in the fourth panel showing an upwards arrow and dollar signs). The task of the participants was to choose which stimulus type (houses or faces) was more associated with receiving reward points through trial-and-error feedback processing. If participants choose the non-set winner (i.e., the one not associated with getting reward points), they received loss feedback (i.e., red box including downwards arrow and dollar sign).

Localizer task: Following the C-PLearn task, participants completed a localizer task, which was originally designed to generate potential regions of interest for assessing the face-responsive sensory cortices in our prior work with adults (van den Berg et al., 2019). We used a very similar localizer task to that one used in our prior work with young adults (van den Berg et al., 2019), which consisted of 20 blocks of 10 trials each. During the task, participants saw single versions (not paired) on each trial of the same house and face images presented during the C-PLearn task, but 20% were somewhat blurred, which served as infrequent targets, while the rest were clear (i.e., non-targets) (see (van den Berg et al., 2019) for more details). The job of participants was to press a button for any blurry images and indicate whether they were a face or a house by pressing a button. But data from the localizer task was not included in our major analysis, as it was not our major questions of interest. Rather, we just collapsed the data over the faces and houses, and focused on other questions in this paper.

### 2.3. Computational modeling of reward learning behavior

To calculate each subject’s reward-learning behavior within the PRL framework, we defined win-stay (WS) trials as those in which the previous trial (*n* − 1) was rewarded (won) and the choice made on the current trial *n* was the same choice as that on trial *n* – 1 (i.e., “stayed” with the same choice of face or house made on the previous trial). We then calculated WS probabilities as the proportion of trials with WS behavior given a previously rewarded trial (i.e., after receiving gain feedback). We defined lose shift (LS) trials as those in which the previous trial (*n* − 1) was not rewarded (i.e., receiving loss feedback) and the choice made on the current trial *n* differed from the choice on trial *n* − 1. We then calculated LS probabilities as a proportion of trials with LS behavior given that the previous trial was unrewarded (i.e., after receiving loss feedback).

### 2.4. EEG Data Collection and Processing

Continuous EEG was recorded using a 128-channel HydroCel Geodesic Sensor Net (Electrical Geodesics, Eugene, OR) and Net Amps 400 series amplifiers at a sampling rate of a 1000 Hertz (Hz). Data was referenced online to the vertex (Cz) during acquisition, and impedances were maintained below 50 kilohms throughout the paradigm.

Offline EEG data preprocessing was performed using EEGLAB version 2021 (Delorme & Makeig, 2004) in MATLAB R2019b (The MathWorks Inc, Natick, MA) using custom scripts (codes are available at: https://github.com/gaffreylab). According to the Maryland analysis of a developmental EEG pipeline recommended for pediatric populations (Debnath et al., 2020), we followed several key preprocessing steps as follows. First, twenty-four channels located on the outer ring of the sensor net were removed from the analyses. As participants were required to wear masks during the EEG sessions according to Duke COVID-related policy, which tended to impair sensor connectivity, these channels had a particularly large number of artifacts during the recordings in these children. The data were then downsampled to 250 Hz, low-pass filtered at 40 Hz, and segments of data without relevant events (i.e., breaks) were removed. Using the ERPLAB v8.2 plugin (Lopez-Calderon & Luck, 2014), a 0.1 to 30 Hz, 4^th^ order Butterworth, bandpass filter was applied. The CleanLine plugin (Kappenman & Luck, 2012) was used to remove any remaining 60 Hz line noise (Mitra & Bokil, 2007). All data were re-referenced to the average of the two mastoids. Bad channels were removed by running the Clean Rawdata plugin: a channel was considered bad if 1) it was flat for more than five seconds, 2) contained more than four standard deviations of line noise relative to its signal, or 3) correlated at less than.8 to nearby channels.

According to current recommendations in the field (Debener et al., 2010; Debnath et al., 2020), a “copy” of the data was made and then, the following steps were applied (Delorme & Makeig, 2004): 1) the high-pass filter at a 1 Hz (4^th^ order Butterworth), 2) Artifact Subspace Reconstruction (ASR;(Mullen et al., 2015) with the burst criterion set to 20 (Chang et al., 2018) to remove large artifacts with Clean Rawdata, and 3) extended infomax Independent Component Analysis (ICA; (Lee et al., 1999) with PCA dimension reduction (50 components). The resulting ICA matrix was then copied over to the original full-length data (i.e., the data just before the copy was made and a 1 Hz high-pass filter was applied). The ICLabel plugin (Pion-Tonachini et al., 2019) automatically removed independent components with a probability greater than.7 of being associated with eye movements and blinks. Application of the ICA matrix to the full-length data and subsequent removal of eye-related components allowed for the preservation of data that would have otherwise been removed by ASR or other artifact removal methods. Bad channels that were previously removed from the analyses were interpolated back into the data set using spherical splines (Perrin, 1989).

For both the C-PLearn task and the localizer task, epochs were extracted from 400 ms before until 800 ms after the onset of the relevant stimulus events (either the choice cue-pairs or the feedback screens in the main task or the single face/house images in the localizer task). All epochs were baseline corrected using the baseline period from –200 ms to the onset of the event. Artifact rejection using the TBT plugin (Trial-By-Trial basis; (Ben-Shachar, 2018)) removed epochs with at least 10 channels meeting the following criteria: 1) peak-to-peak amplitudes exceeding 100 μV within 200 ms windows sliding across the epoch by 20 ms increments, 2) voltages below –150 uV or greater than +150 μV, or 3) joint probabilities above three standard deviations for local and global thresholds. If less than 10 channels in an epoch met criteria for rejection, the epoch was not removed, but the identified channels were interpolated for that epoch only. However, if there were more than 10 channels in the epoch that met criteria for rejection, the epoch was not included. Lastly, the averaged ERPs for each bin (i.e., Gain, Loss) for each subject were computed. Mean of the number of accepted Gain and Loss trials per subject were 184.3 and 128.2, respectively.

### 2.5. Data binning and averaging

From the C-PLearn runs, the epoched EEG data were binned according to feedback (gains and losses) and choice (face and house), which resulted in an average number of trials for each subject (Brown et al.) for face gain [90.53 (± 24.97)], face loss [62.13(±13.73)], house gain [93.76(±19.60)], and house loss [66.06 (±14.76)], after rejection of noisy epochs. As expected, due to learning, there were significantly more gain trials than loss trials [*F*(1,29) = 43.52, *p* <.001]. On the other hand, there were no significant differences in the number of gain or loss trials for choosing a face vs a house ([*F*(1,29) = 0.46, *p* =.50].

### 2.6. Analysis of cue-evoked ERP data

We calculated the N2pc response evoked by the choice cue-pair at the beginning of each trial to assess attentional orientation to the two stimulus types as a function of what they will later choose that trial, following a similar procedure used in our prior work in adults (van den Berg et al., 2019). More specifically, the N2pc neural responses were assayed by the standard N2pc contralateral-vs-ipsilateral analysis this component (Luck & Kappenman, 2012), that is by subtracting the activity in the contralateral channels (relative to the chosen side) minus the ipsilateral channels and then collapsing over the left and right sides. Further, we also calculated the N2pc amplitudes evoked by the choice-cue responses as a function of whether the cue was a set-winner or not.

### 2.7. Statistical Analysis

For statistical analysis, we first calculated the time-locked ERPs as a function of the various event types and conditions. Based on previous literature, the reward-related Positivity (RewP) was measured from 275 to 375 ms following the feedback stimulus in a fronto-central ROI (E10, E11, E16). The P300 was measured from 400 to 600 ms following the feedback stimulus in a central-parietal ROI (Left/Right: E37/E87, E31/E80, E53/E86, E60/E85, E67/E77, E61/E78, E54/E79; middle line: E129, E55, E62, E71; (Fischer & Ullsperger, 2013; San Martín, 2012; San Martín et al., 2013). Subsequently, the extracted values were analyzed with a repeated-measures analyses of variance (ANOVAs) to test for statistical significance (*p* <.05) using the SPSS 2021 version.

The cue-related attentional bias (N2pc) was measured from 175 to 225 ms after onset of the choice-cue image pair, from corresponding left and right occipital ROIs (E50/E101, E58/96; (Hickey et al., 2010; Kappenman & Luck, 2012; San Martín et al., 2016)). Subsequently, we calculated the difference in voltage contralateral vs ipsilateral relative to the side on which the set-winner was presented, as well as contralateral vs. ipsilatateral to the side the stimulus that would be chosen on that trial.

The EEG data collected during the independent localizer task were used to confirm that the stimuli that we used in this study elicited face-vs-house activity differences as has been observed in the existing literature and to delineate potential influences of reward learning on stimulus-specific regions. To do so, we first visually inspected all posterior electrodes (i.e., over visual cortex) for the ERP differences between the neural responses to face and house images to make sure these effects were representative of previously reported findings with adults, including the hallmark, face-selective, lateral inferior occipital N170 responses. The N170 was measured from 170 to 220 ms after onset of the cue image, from corresponding left and right occipital ROIs (E66/E84, E67/E91, E61/E92).

To calculate each subject’s individual behavioral learning rate, we used the same formula as in van den Berg et al. (2019) and calculated on a trial-by-trial number basis using a mixed-modeling approach using the *lme4* statistical package (Bates et al., 2015). A varying slope of condition per subject (the random effect) was included in the model if the Akaike Information Criterion (a measure of the quality of the model) improved. Statistical significance was set at *p* < 0.05, with the Satterthwaite’s degrees of freedom method as given by the *R* package *lmerTest* (Kuznetsova et al., 2017).

## 3. Results

### 3.1. Behavioral Results

Consistent with our prior work in adults (van den Berg et al., 2019), and as predicted, behavioral measures in the children demonstrated their reinforcement learning ability based on trial-by-trial reward-related feedback. According to a *post-hoc* paired *t*-test, the percent of children choosing the block winner on the last (18^th)^ trial of the block (*M* ± *SD*: 0.75 ± 0.16) was significantly higher compared to on the first trial (*n*=30, *M* ± *SD*: 0.42 ± 0.10) (*t* (29) = –9.54, *p* <.001)). As presented in Figure 2.a, at the beginning of each 18-trial set, participants chose the most likely winner for that set at only chance level (0.45-0.50), as would be expected. But by the end of the set, this proportion of choosing set-winner increased up to ∼0.75 on average, indicating that the children had used the feedback across the trials of that set to learn to choose the more likely probability-based winner. Nevertheless, the individual learning curves revealed very large individual difference in reward learning, as the final proportion of choosing set winners in a particular block ranged from 0.26 to 1.0 across subjects and blocks. Due to this large range, we wanted to explore whether ‘good’ vs. ‘bad’ learners might show different patterns of reward-related ERP components, in particular by dividing the subjects using a median split based on their learning rate (Fig2).

**Figure 2.**
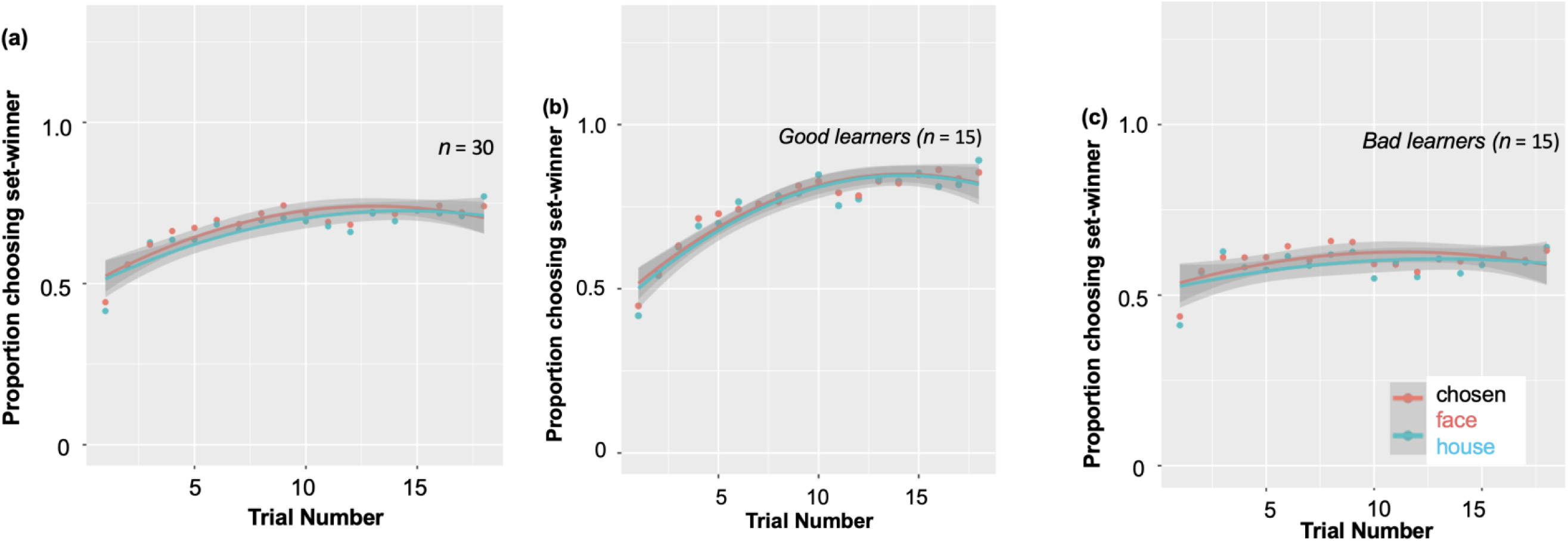
Individual differences in behavioral probabilistic reward learning. During 18 blocks, the proportion of choosing set-winner for all participants ranged from 0.40 to 0.73. The individual learning curves revealed very large individual difference in reward learning, as the final proportion of choosing set winners in a particular block ranged from 0.26 to 1.0. Due to this large individual difference observed, we divided participants into good vs. bad learners according to a median split.

Specifically, as presented in Fig 2a, learning rates for good learners showed continuous increases from the first trial (learning rate: 0.43) through the 9^th^ trial (learning rate: 0.80) (*t*(14) = – 9.57, *p* < .001), to the last trial (learning rate: 0.87) (*t*(14) = 02.27, *p* =.03). On the other hand, learning rates for bad learners showed somewhat increase from the first trial (learning rate: 0.42) to the 9^th^ trial (learning rate: 0.63) (*t*(14)= –3.77, *p* =.002), but no increase during the last half trials (*t*(14)= –0.02, *p* =.98).

As presented in Figure 3, good vs. bad learners adopted different behavior strategies during the C-PLearn task. Good learners were more likely to choose the same stimulus type (i.e., Win-Stay, WS behavior strategy) on the trial after one with a gain feedback (*t*(28) =3.92, *p* <.001, Cohen’s *d* =1.43). In contrast, bad learners were more likely to change their response choice (i.e., Lose-Shift, LS behavior strategy) on the next trial after loss feedbacks (*t*(28) = –2.89, *p* = 0.007, Cohen’s *d* =1.05).

**Figure 3.**
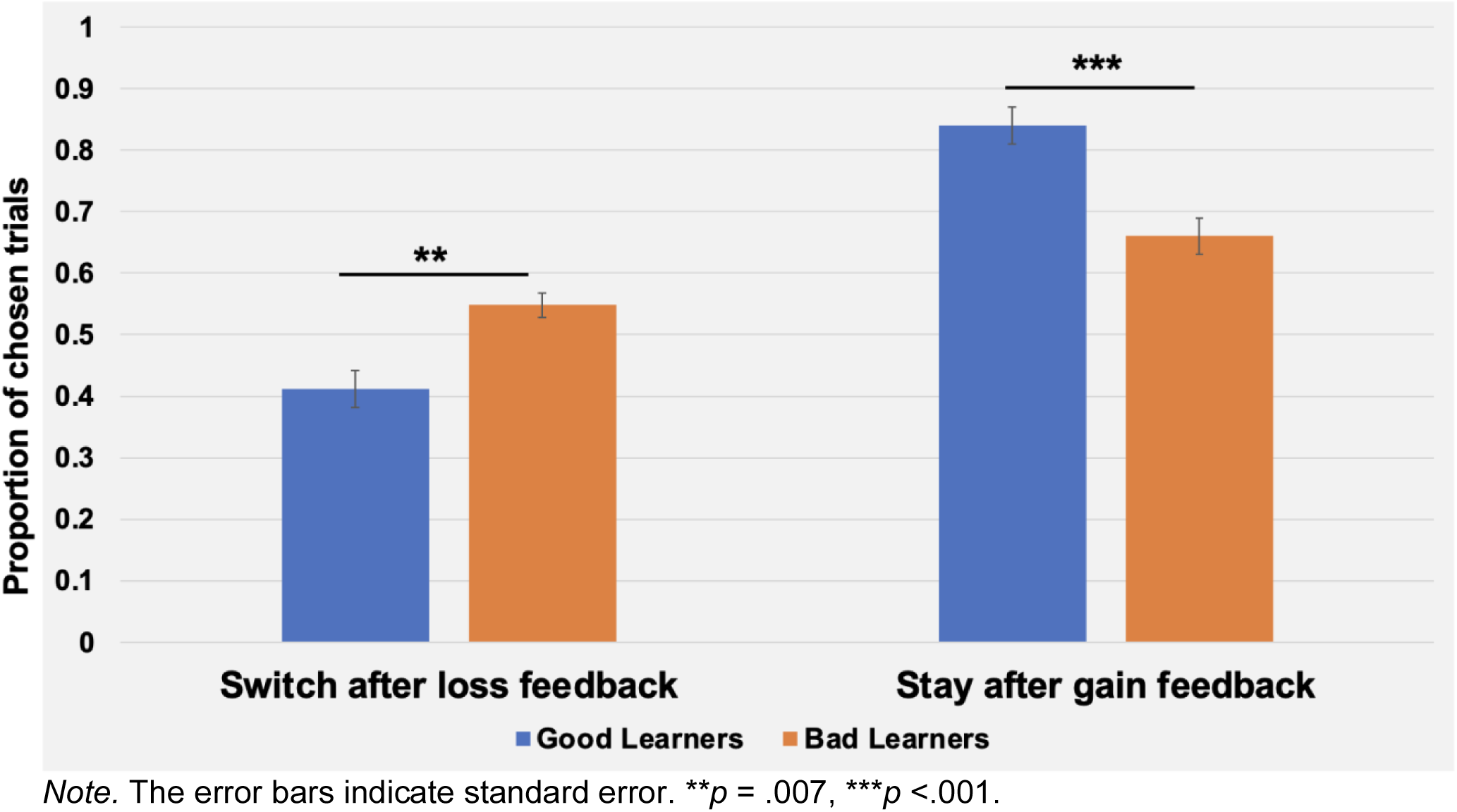
Behavioral strategies during the Child-version of Probabilistic Reward Learning (C-PLearn). Bad learners with low learning rates were more likely to change their choice on the next trial following loss feedback (i.e., Lose-Shift strategy). On the other hand, good learners with high learning rates were more like to make the same choice on the next trial following gain feedback (i.e., Win-Stay strategy).

### 3.2. Electrophysiological Measures

#### 3.2.1. The Reward-related Positivity (RewP)

Consistent with prior work in adults, we anticipated the processing of gains vs. loss feedback would be reflected by the reward positivity wave, that is, the RewP in the fronto-central ROIs. As presented in Figure 4.1, we found the presence of RewP around 275-375 ms after gain vs. loss feedback in our frontal ROI (E10, E11, E16) for young children. Given large individual differences in learning rates in our sample, we compared the RewP amplitudes between good vs. bad learners subgroups. Differing from our expectation, the presence of the RewP amplitudes appear to be mostly driven by bad learners with lower learning rates. The independent *t*-tests revealed that bad learners had greater RewP amplitudes in the frontal ROIs compared to good learners [*t*(27)= –2.14, *p* = .04, Cohen’s *d* = 0.79] (see Figure 4.2).

**Figure 4.1.**
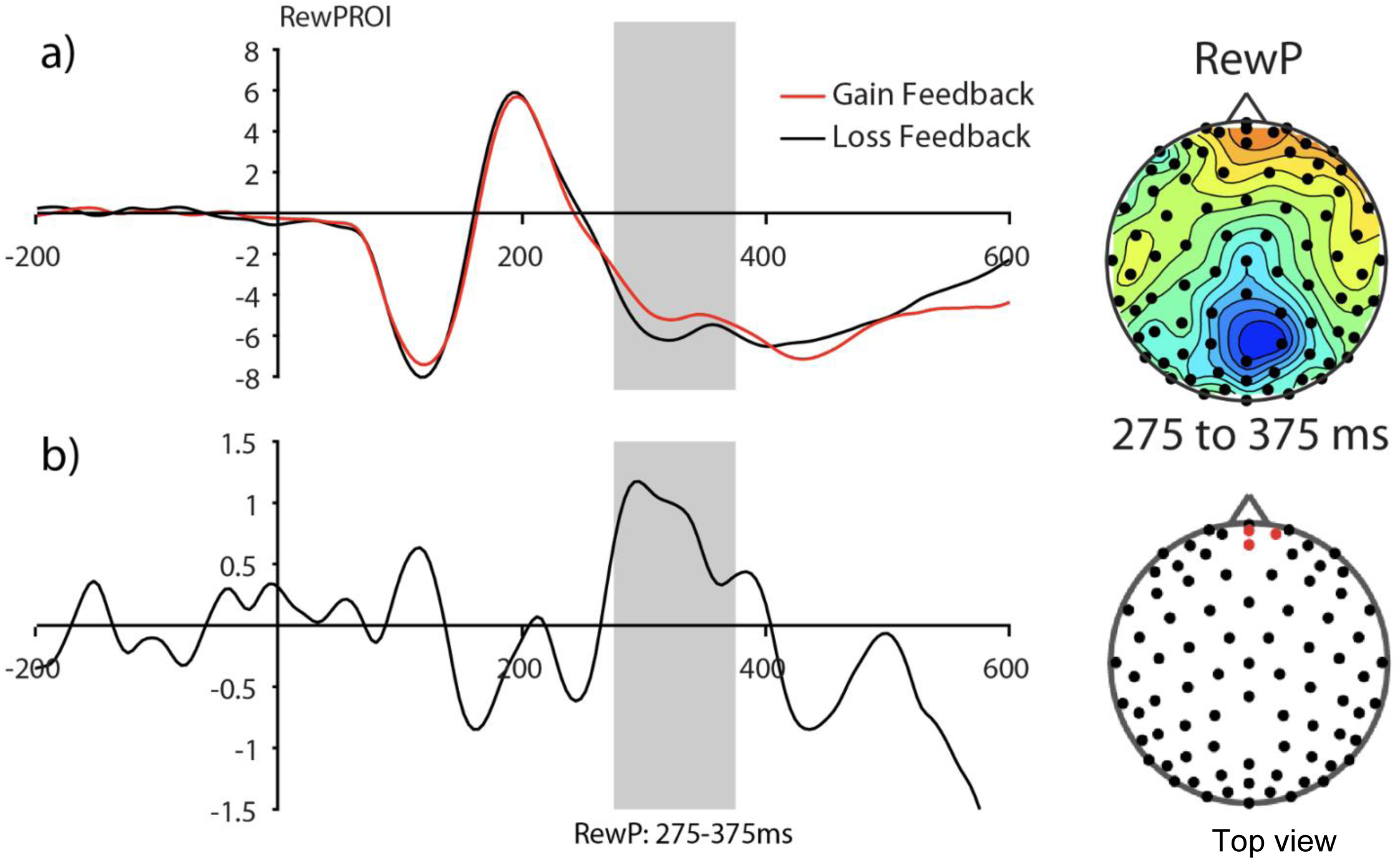
The Frontal reward-related potential (gain vs. loss feedback) in preadolescents (n=30). (a) Grand average feedback-cue evoked responses for Gain and Loss reward feedback from the fronto-central ROI (E10, E11, E16) (see red dots in the topography for the location of the ROI). (b) The reward-related potential difference waveform during Gain minus Loss reward feedback. The reward-related potential was measured from 275 to 375 ms in a fronto-central ROI.

**Figure 4.2.**
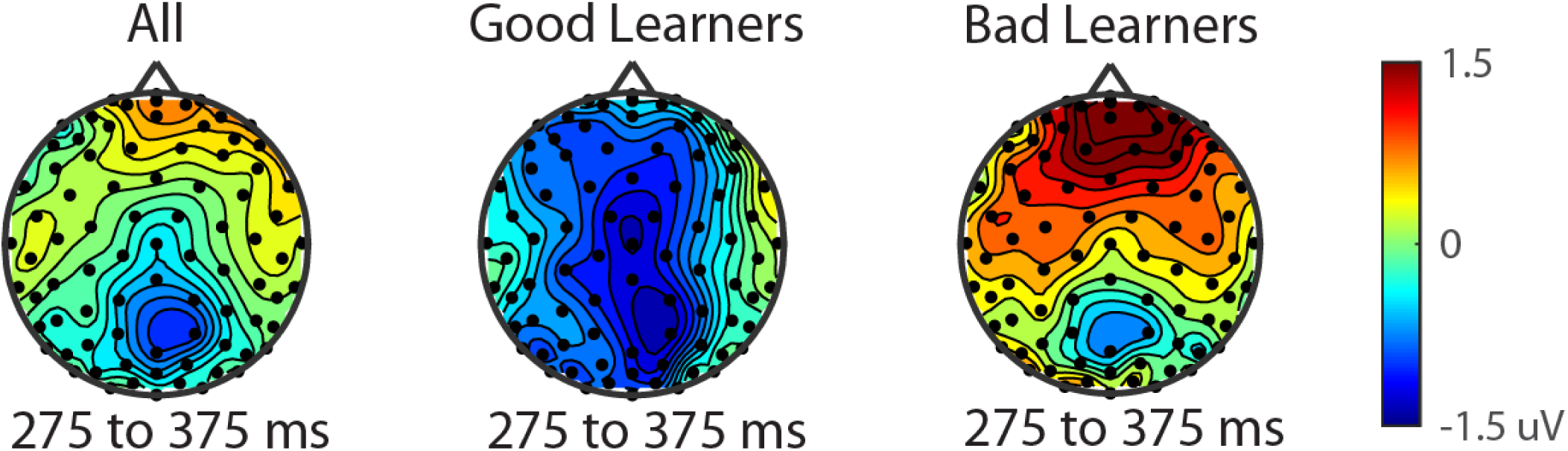
The reward-related positivity between good and bad learners. The comparison of the reward-related potential ERP marker measured from 275 to 375ms in the fronto-central ROI between good and bad learners. Dark red colored electrode sites in the frontal ROI shows significantly greater reward-related positivity amplitudes for bad learners (i.e., those with low learning rates) compared to good learners (i.e., those with high learning rates).

#### 3.2.2. P300 during reward gain vs. loss feedback as a function of behavior strategy

We examined updating reward values as a function of reward learning, as reflected by behaviorally dependent modulation of the P300. Figure 5.1 shows the P300 in response to gain vs. loss feedback as a function of behavior strategy (i.e., Switch vs. Stay) between good and bad learners. Using P300 amplitudes in the central ROIs between 400 and 600 ms for good and bad learners, we conducted repeated measures ANOVAs with two within-subject factors (i.e., behavior strategy: switch vs stay, feedback: gain vs. loss) and one between-subject factor (i.e., good vs. bad learners).

**Figure 5.1.**
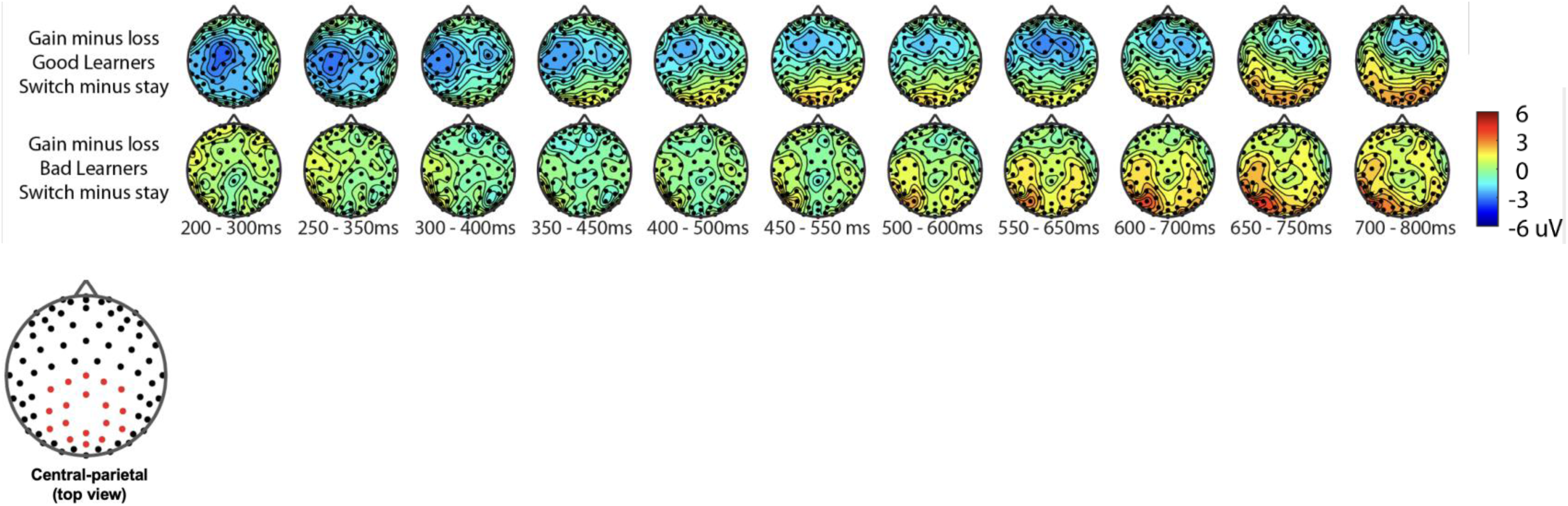
Topography of P300 during gain vs. loss feedback as a function of behavioral strategy. P300 amplitude differences in central and parietal ROIs on trials on which participants switched their responses on the next trial after gain feedback vs. after loss feedback. The red dot in the topography at the bottom shows the location of the central-parietal ROI. Top row: warm colored electrode sites indicate bigger P300 amplitudes on stay trials for good learners with high learning rates. The Bottom row: warm colored electrode sites indicate bigger P300 on stay trials for bad learners with low learning rates.

The ANOVAs were conducted to examine whether bad learners, relative to good learners, showed greater P300 amplitudes in response to gain vs. loss feedback as a function of switch behavior strategy. As shown in Figure 5.2(a), bad learners (i.e., those with low learning rates), relative to good learners, showed higher P300 amplitude differences in central, parietal ROIs on trials when they switched their responses on the next trial after gain feedback vs. after loss feedback. The repeated ANOVA analysis revealed no main effect on the P300 of learner type [bad vs. good: *F*(1, 27) = 1.45, *p* =.23, 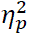: 0.05]. We did observe, however, main effects of behavior strategy with the greater P300 amplitudes on lose-switch trials than win-stay trials [*F*(1, 27) = 9.11, *p* =.005, 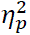: 0.25] and feedback with greater P300 amplitudes on loss compared to gain trials [*F*(1, 27) = 7.00, *p* =.01, 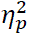: 0.21], as presented in Figure 5.2.(b) and (c), respectively, but no interaction of feedback by behavior strategy [*F*(1, 27) = 0.34, *p* =.56, 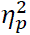: 0.01] or of behavior strategy by learner type [*F*(1, 27) = 0.38, *p* =.54, 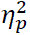: 0.01] or feedback by behavior strategy by learner type [*F*(1, 27) = 0.09, *p* =.76, 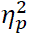: 0.003] (see Figure 5.2). According to a *post-hoc* paired *t*-test, however, after receiving gain vs. loss feedback, children showed greater P300 amplitude differences in the central, parietal ROIs when they switched their choice on the subsequent trial compared to when they chose the same response [*t*(28)= 2.68, *p* = .012, Cohen’s *d* =0.32] (see Figure 5.2.b). Also, P300 amplitudes in the same central and parietal ROIs were higher when children received loss feedback compared to gain feedback, collapsed across both good and bad learners [*t*(28)= 3.03, *p* = .005, Cohen’s *d* = 0.39] (see Figure 5.2.c), consistent with adult findings (van den Berg et al., 2019).

**Figure 5.2.**
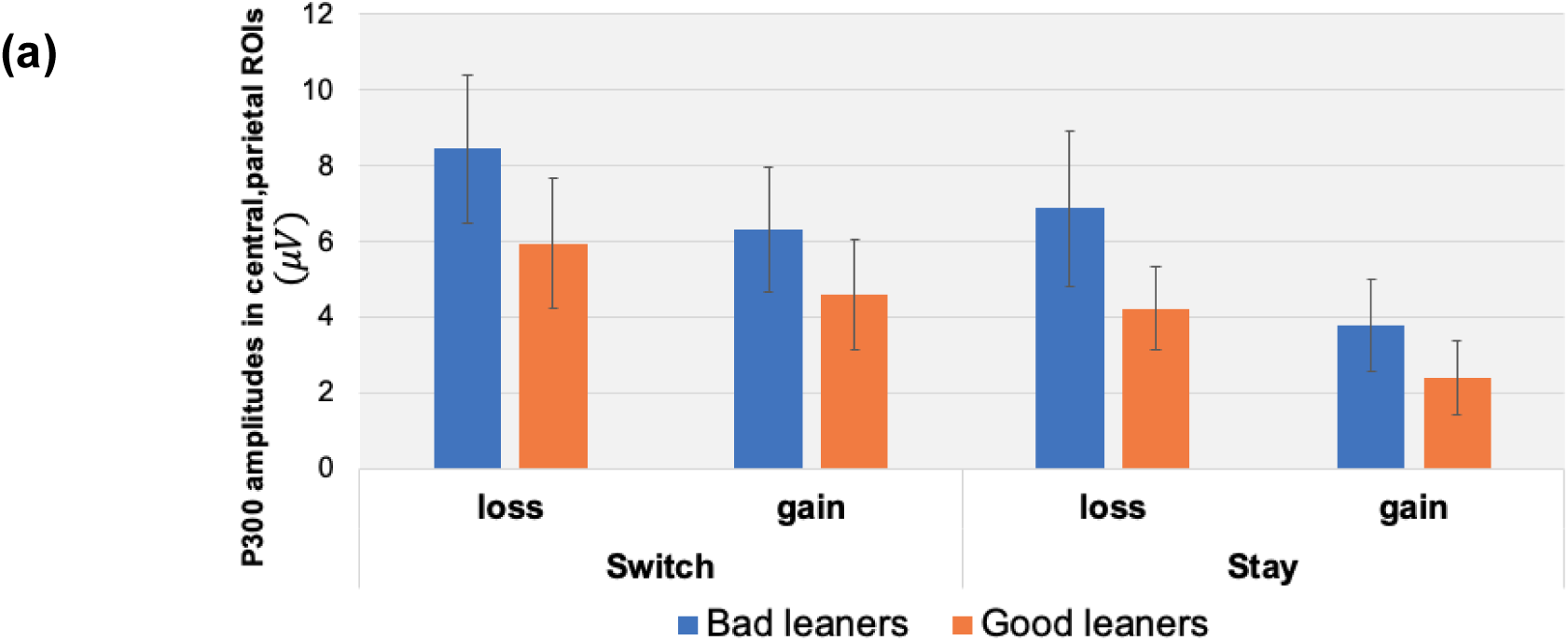
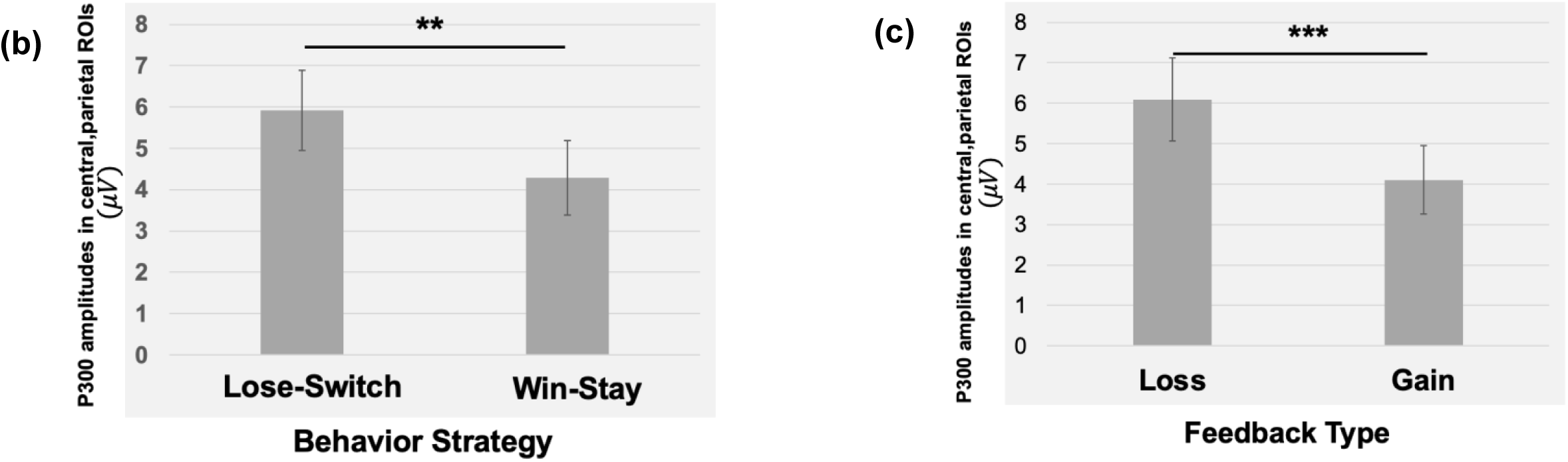
P300 amplitudes in central, parietal ROIs as a function of feedback type and behavioral strategy. (a) Using P300 amplitudes in the central –parietal ROIs between 400 and 600 ms for good and bad learners, we found that bad learners (i.e., those with low learning rates), relative to good learners, showed higher P300 amplitude differences in central, parietal ROIs when they switched their responses on the next trial after gain feedback vs. after loss feedback. (b) For all children, we found greater P300 amplitudes in the central-parietal ROI when participants switched their choice on the next trial after loss feedback (i.e., lose-switch strategy) compared to when they chose the same choice after gain feedback (i.e., win-stay strategy). (c) The P300 amplitudes in the same central-parietal ROIs were higher when participants received loss feedback compared to gain feedback. *Note.* Error bars indicates standard error; asterisks indicate significance level: \*\**p* = 0.01, \*\*\**p* =.005.

**Figure 6.**
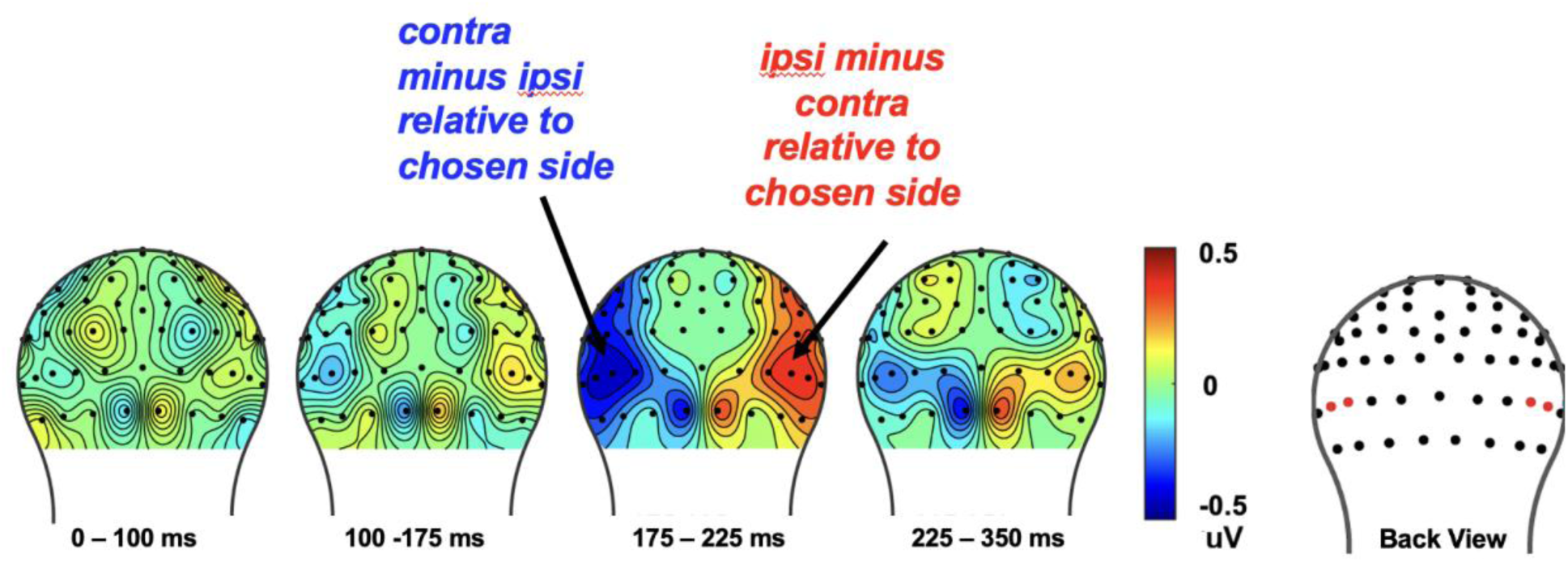
The cue-evoked attentional bias of N2pc towards the to-be-chosen side on each trial. To increase signal-to-noise ratio, we collapsed over whether the chosen image was that of a face or a house and analyzed the effect of choice behavior on the N2pc over the course of the 18-trial set. The cue-evoked N2pc amplitudes measured after onset of the cue image pair, from corresponding left and right occipital ROIs. was found from 175 to 225ms (see red dots in the topography for the location of the ROI electrodes)

#### 3.2.4. N2pc reflects attentional bias towards choice behavior

In our prior work in college students (van den Berg et al., 2019) we found that the N2pc changed across the block for the set winner as that set winner was learned. In the current study, however, we did not find such an effect for the set winner, but only for the behavioral choice regardless of whether or not reward learning had been consolidated. First, we analyzed the effect of choice on the N2pc over the course of the 18-trial set by collapsing over whether the chosen image was that of a face or a house (to increase the signal-to-noise ratio). Then, we analyzed the relationship between the N2pc elicited by the cue-pair stimulus relative to the behavioral choice made later in the trial, regardless of what the set-winner was. As expected, we found the presence of an N2pc for the to-be-chosen stimulus type in the latency of 175-225 ms after the onset of each cue-pair stimulus (E50/E58) [*t*(29)= –2.85, *p* <.008]. These results indicate that children shifted their attention to the stimulus type that they were going to choose that trial. Additional independent *t*-tests on the N2pc amplitudes as a function of the set winner, measured in the ROIs from 175 to 275 ms between good and bad learners, revealed no statistical differences (*t*(28) = –1.76, *p*=.09), even though the behavior indicated that the good learners were learning the reward associations better.

## 4. Discussion

Consistent with our prior work in adults (van den Berg et al., 2019), our behavioral C-PLearn findings indicate that, at least at a group level, preadolescents are able to learn which stimulus type in a pair (a face vs. a house) is more likely to result in a higher probability of reward over a series of trials. However, we observed large individual differences in individual learning rates and divided the participants into good learners (i.e., those with high learning rates) vs. bad learners (i.e., those with low learning rates). This revealed that bad learners tend to use a different behavioral strategy than did good learners, namely a “Lose-Shift” behavior strategy rather than a “Win-Stay” strategy. In line with prior work with adults (Worthy & Maddox, 2014), choosing the same choice on the next trial after receiving reward feedback –-namely, a “Win-Stay” strategy –-appears to be the better heuristic learning strategy in these binary decision-making situations, including in preadolescent children.

At the neural level, the current study’s findings support the C-PLearn task as a developmentally appropriate one for identifying and measuring key electrical brain activations associated with PRL processes and behavior in preadolescents. In contrast to our predictions, we found that poor preadolescent learners with low learning rates showed greater amplitudes of the RewP ERP components than did good learners, indicating greater reward-feedback sensitivity. According to the prominent RL theory of the RewP (or the sometimes studied inverse measure of the feedback-related negativity), this reward-sensitive process is supported by the dopamine system of the brain (Holroyd & Coles, <u>2002</u>). This theory is based on many animal and human studies implicating the basal ganglia and midbrain dopamine system in reward prediction and RL, with these processes modulating cortical activations that are reflected by in the fronto-central RewP ERP in humans. Briefly, the basal ganglia evaluate ongoing events in the environment and predict whether the events will more likely lead to success or failure. When the basal ganglia revise the predictions for the better, they induce a phasic increase in the activity of midbrain dopaminergic neurons. But, once the stimulus-reward outcome associations have been learned, there is a phasic decrease in the activity of midbrain dopaminergic neurons during reward delivery but a phasic increase in the activity of midbrain dopaminergic neurons during the presentation of reward-predictive cues (reviewed in (Daw & Tobler, 2014). For example, according to seminal animal work by (Schultz et al., 1997)), when a reward was consistently paired with a predictive stimulus the phasic increase in dopamine firing rate observed at the time of reward delivery *diminished over time* and instead a phasic increase in dopamine firing rate was observed shortly after the onset of the predictive stimulus. More recently, this has also been demonstrated in humans. According to the Krigolson et al. (2014) study with adults (n=18), novel rewards elicited neural responses at the time of reward outcomes that *rapidly decreased in amplitude with learning*. In line with this RL framework (Holroyd & Coles, 2002), we speculate that the blunted RewP amplitudes observed in good learners with high learning rates during reward outcomes might reflect such phasic decrease in dopamine firing rate with learning, but they might have shown the RewP increases at the reward-predicting cues. In contrast, poor learners with low learning rates might have showed greater RewP during the reward outcome period, as they didn’t learn the stimulus-outcome associations. However, this speculation needs to be more fully tested in future studies with larger numbers of subjects.

We also observed indication of rapid spatial attention modulation toward the stimulus to be chosen regardless of the set-winner across blocks, as evidenced by the N2pc, as we had observed previously in young adult subjects in a similar paradigm (van den Berg et al., 2019). However, we did not find an increasing N2pc toward the set winner stimulus type that was learned across a block, an effect that we had observed in young adults (van den Berg et al., 2019). Lastly, our preadolescent group showed contextual modulation of the feedback-locked P300 over the parietal cortex as a function of behavior strategy (i.e., Win-Stay-Lose-Shift). Thus, these results provide novel insights into the associations between RL-related ERP components and computationally-driven reward learning behavioral strategies (Hernstein et al., 2000). Accordingly, the current study supports the view that the C-PLearn task can be very useful for eliciting key measurable behavioral and neural correlates of PRL processing in preadolescent children.

To our knowledge, this is the first study to not just examine the key neural processes underlying PRL learning (i.e., reward-related feedback processing and attention orienting) in preadolescents, but also with linkage to a combination of PRL and WSLS models. Most existing ERP studies with children have focused mainly on the RewP effect on general reward feedback processing (e.g., (Burani et al., 2019; Ethridge et al., 2017; Ferdinand et al., 2016; Kessel et al., 2016; Kessel et al., 2019; Lukie et al., 2014). However, little is known about the RewP in the context of reinforcement learning in preadolescents. This is a critical gap of knowledge in the field because poor PRL learning may be a key factor underlying adolescent-onset of depression (Chen et al., 2015; O’Callaghan & Stringaris, 2019). Although such mechanisms are not yet clear, a recent meta-analytic review presented strong evidence that depression is associated with disordered brain signals of reward prediction error and with poor reinforcement learning behavior (Halahakoon et al., 2020). Here we extend such research by examining the RewP effect in the domain of RL processing.

Given that we observed large individual differences in computationally-driven behavioral-learning rates in the present study, we compared the RewP effect between “good” and “bad” learners, as defined by their computationally-derived learning rates. This adaptation of the WSLS strategy model in the current study revealed that poor preadolescent learners tended to more automatically “switch” their choice following losing feedback, a finding consistent with prior work with college students (Worthy & Maddox, 2014). The WSLS is a rule-based strategy that has been shown to be commonly used in binary outcome choice tasks (e.g., Otto et al., 2011). According to Worthy and Maddox (2014), college students with higher probabilities of shifting following a loss trial (i.e., “Lose-Shift” behavior strategy) showed poorer learning performance. Along with prior work with adults, our current findings add converging evidence that combining the WSLS and PRL models may provide a better account of decision-making behavior even in preadolescents compared to a single PRL model. Also, it is possible that preadolescents with greater reward sensitivity, as reflected by greater RewP amplitude, might have been more distracted when exposed to cues that were previously associated with high reward values but were associated with non-reward outcomes. Such distraction might have led to choosing inefficient learning strategies, such as the Lose-Shift strategy. Indeed, a recent meta-analytic review by (Rusz et al., 2020) showed converging evidence for strong reward-driven distraction effect on reward-learning mechanisms. For example, college students (mean age = 22.3 years) were less accurate when they were exposed to irrelevant cues that had previously been associated with high reward value (Rusz et al., 2018).

One key component of PRL processing is the instantiation of attentional shifts towards stimuli associated with higher probability of getting rewards. Previously learned reward values can have a pronounced effect on the allocation of selective attention, which can be indexed in the present paradigm by an increasing amplitude of the attention-sensitive, lateralized negative deflection (the N2pc) contralateral to the set-winner in the cue-pair presentation. The prior ERP studies in adults showed that once a stimulus-reward association has been learned, it tended to induce a larger attentional shift when presented, while also predicting the likelihood of the choice to be made on that trial (Hickey et al., 2010; Hickey & van Zoest, 2012; San Martín et al., 2016; van den Berg et al., 2019). Such pronounced attentional shift in the context of reward learning, as indexed by the N2pc, has been little investigated in children. Only a handful of ERP studies with children reported the role of N2pc during visual search (Couperus & Quirk, 2015; Li et al., 2022; Shimi et al., 2014; Sun et al., 2018; Turoman et al., 2021) and/or in working memory tasks (Rodríguez-Martínez et al., 2021; Shimi et al., 2015). In addition to the RewP effect, the current study revealed the modulation of N2pc ERP amplitude indexing rapid attentional orienting towards the choice behavior (i.e., the stimulus that would be chosen on that trial). Our prior work in adults (van den Berg et al., 2019) revealed the presence of significant N2pc to the to-be-chosen stimulus *<u>and</u>* to the more rewarded stimulus in a set, particularly on the latter trials of the set after more learning of the set winner was able to occur, indicating rapid attention modulation as a function of reward learning. Here, in preadolescents, we found a modulation of attention bias towards the to-be-chosen side, which was elicited similarly over the whole course of learning, but we were not able to replicate the influence of reward-learning on the rapid attention modulation towards the more rewarded stimulus. We speculate any learning-based modulation of the attention-sensitive N2pc amplitudes may have been too small to be observed here, because it was occurring only in the good learners and only during the later trials of the blocks. This speculation should be tested in future studies with larger samples of both good and bad child learners.

Although we found that, like college students, preadolescents learned the stimulus-reward outcome association at a group level, there were large individual differences in behavioral learning rates. With the computationally-driven PRL framework, our behavior data indicated major differences in strategy between good and bad learners, with good learners using win-stay strategy and bad learners using more of a lose-switch strategy. PRL theory generally assumes that the most recent outcomes exert the most influence on the current choice, such that behavior on a given trial, *n*, depends particularly upon the choice and outcome on the preceding trial, *n* − 1. This relationship has been formalized as a simple, heuristic learning strategy called Win-Stay-Lose-Shift (WSLS: (Hernstein et al., 2000). In our prior work in college students (San Martín et al., 2013), the amplitude of the feedback-locked P300 over central and parietal cortex, but not of the FN (RewP), predicted behavioral adjustments (i.e., lose-switching strategy) on the next trial. These results are in line with the context-updating hypothesis of the P300 ERP component (Nieuwenhuis et al., 2005): accordingly to this view, decisions on each trial are informed by an internal model of the symbol/probability(win) contingencies, and the P300 amplitude reflects the extent of the feedback-triggered revision of such a model. Thus, the feedback-locked P300 distributed over central and parietal cortex may reflect adjustment processes to maximize gain and/or minimize loss in the future. In line with this view, the present study also found a modulation of P300 over central and parietal cortex as a function of behavioral strategy and feedback type, in that the feedback-locked P300 amplitudes were greater on lose-switching trials compared to win-staying trials, and in response to loss vs. gain reward feedback.

In contrast to our expectation, however, we did not find overall differences in the feedback-locked P300 amplitudes located over central and parietal cortex between good and bad learners, even though our data indicated a modulation of the P300 amplitudes as a function of behavioral strategy and feedback more generally. We cannot rule out a possibility that there was not enough power to detect the interaction effect of P300 amplitudes and subgroup types due to limited number of participants.

The greatest limitation of the current study is having a relatively small sample size for examining individual level differences and differential subgroup interactions, which may have resulted in our missing more nuanced distinctions in our behavioral measures of RL behavior and brain function (e.g., no significant N2pc to the stimulus with a higher likelihood of reward – i.e., the set winner – as that reward association was being learned). In addition, our sample was limited to preadolescent children. Accordingly, future work will be needed to replicate or extend the current findings and test their generalizability in other age ranges and in larger groups reflective of the general US population in terms of race and ethnicity.

In summary, the results of the current study provide novel insights into how a cascade of core neurocognitive processes underlying RL are associated with preadolescents’ behavior learning strategy: 1) The RewP component was modulated by individual learning rates in preadolescents: good learners with high learning rate appear to show more blunted RewP amplitudes with their learning; 2) There was an early attentional orienting towards the stimulus that would be chosen on a trial, as evidenced by the N2pc, but such orienting processes were not observed as a function of the learning of the reward associations; and 3) There was evidence of an updating reward values as a function of behavior strategy, reflected by the modulation of P300 amplitudes depending on lose-switch vs. win-stay strategy in the different learning groups. The alterations in these cognitive processes have been previously associated with the likelihood of adolescent-onset depression (Chen et al., 2015). More specifically, adolescents have showed heightened vulnerability for depression with the onset of puberty when there are differences in RL behavior, suggesting a differential organization or development of frontal-striatal reward brain circuitry (Kessler et al., 2001; Walker et al., 2017). The current study shows that the CPLearn task with the use of EEG measures of brain activity may be a promising tool to further investigate the relationships between attentional processing, learning, and reward sensitivity in preadolescents, which is turn may advance our understanding of the factors underpinning the risk for depression and other cognition-development issues in this key demographic group.

## 1. Author Contributions

Chung Y.S. led all aspects of the study, including the development of the task paradigm, the collection of the data, the analyses of the data, the interpretation of the results, and the writing of the manuscript. Van den Berg B. and Roberts K.C. contributed to the development of the C-PLearn task, the preprocessing pipelines, and the data analyses. Bagdasarov A. contributed to development of preprocessing pipelines. Woldorff M.G. contributed to the development of the task paradigm, provided guidance on the EEG data analyses, and contributed to the interpretation of the results and feedback on the manuscript. Gaffrey MS. contributed to the development of the task paradigm, guidance on the analyses, interpretation of the results, and feedback on the manuscript, and provided the research funding to support the study.

## 3. Acknowledgments

We gratefully acknowledge the children and their parents who participated and completed this study despite challenges due to the COVID pandemic (i.e., wearing face masks throughout all sessions).

Also, we would like to specially thank to research staff Amanda Neal and Summer Lawrence for their effort and support of data collection. Without their help, we could not have successfully completed this project.

This paper was posted online in preprint (Chung et al., 2023) before being submitted.

## 4. Supplementary Material

None.

## 5. Data Availability Statement

The datasets and coding used in this study are available upon request.

